# Traction force dynamics during patient-derived glioblastoma neurosphere invasion in 3D Matrigel

**DOI:** 10.1101/2024.10.01.616178

**Authors:** Hyeje C. Sumajit, David Böhringer, Falak Syed, Gengqiang Xie, Taylor Gogolen, Vidhisha Gautam, Shiv Patel, Branko Stefanovic, Yan Li, Christoph Mark, Jerome Irianto

**Affiliations:** Department of Biomedical Sciences, College of Medicine, Florida State University, Tallahassee, FL, USA; Friedrich-Alexander University Erlangen-Nürnberg, Department of Physics, Erlangen, Germany; Department of Chemical and Biomedical Engineering, FAMU-FSU College of Engineering, Florida State University, Tallahassee, FL, USA

## Abstract

**Purpose:** Glioblastoma multiforme (GBM) is an aggressive brain tumor with a 5-year survival rate below 7%. Poor outcomes are driven in part by diffuse invasion into surrounding brain tissue, which limits complete surgical resection and promotes recurrence. Improved therapies require a quantitative understanding of the mechanics of GBM invasion.

**Methods:** Patient-derived GBM neurospheres were embedded in Matrigel and analyzed using live imaging, three-dimensional traction force microscopy (3D TFM), and targeted cytoskeletal perturbations.

**Results:** Invading GBM cells adopted an elongated, protrusion-rich morphology, with F-actin enriched at the cell periphery and microtubules extending along the protrusion shaft, consistent with a mesenchymal-like invasion program. 3D TFM revealed sustained matrix engagement, including progressive bead clustering, increasing cumulative traction forces, and traction hotspots concentrated near protrusion tips. Perturbation studies separated cytoskeletal requirements for invasion and force transmission: actin polymerization was essential for invasion, myosin II activity was required for robust traction generation and efficient invasion, and microtubule polymerization supported directional persistence and maximal traction output. Notably, a low level of invasion persisted under myosin II inhibition despite minimal detectable traction forces, and this residual invasion was not suppressed by pan-MMP inhibition, indicating a traction-poor, MMP-independent invasion component in Matrigel.

**Conclusions:** These findings establish a quantitative mechanical framework for GBM neurosphere invasion in 3D Matrigel and define distinct contributions of actomyosin contractility and microtubules to invasive progression.

## Introduction

Glioblastoma multiforme (GBM) is among the most aggressive and invasive brain tumors, with a 5-year survival rate below 7%^1^. Unlike many solid tumors, GBM rarely metastasizes outside the brain but instead disseminates locally through the perivascular space and brain parenchyma, producing diffuse margins that preclude complete surgical resection and drive recurrence^2-5^. Defining the cellular mechanisms that enable invasion in 3D microenvironments is therefore central to improving therapeutic strategies for GBM.

Tumor cells can switch between migration programs as they navigate complex extracellular structures, engaging distinct adhesion, cytoskeletal, and signaling modules in response to matrix architecture and confinement^6-8^. Rho-family GTPase pathways regulate actin organization and activate non-muscle myosin II, enabling cellular traction force generation, morphological change, and invasion through dense extracellular spaces^9^. Similarly, in GBM, myosin II activity facilitates deformation of the cell body and nucleus, supporting translocation through narrow interstitial spaces^10,11^. Invasive GBM cells often extend elongated protrusions at the leading edge, followed by actomyosin-driven contraction that advances the cell body and retracts the rear^10,12^. Indeed, myosin IIA inhibition suppresses invasion in established GBM models^13^, suggesting the critical role of myosin II activity in GBM invasion.

Here, patient-derived GBM neurospheres embedded in Matrigel were used as a 3D invasion model to couple invasion dynamics with quantitative measurements of cell-generated forces. Live imaging and confocal microscopy were used to resolve invasion morphology and cytoskeletal organization, and 3D TFM was applied to reconstruct matrix deformations and traction forces during invasion. Using pharmacologic inhibition of actin polymerization, microtubule polymerization, and myosin II activity, the contributions of cytoskeletal architecture and contractility to invasion and force generation were tested. The data support a prominent role for actomyosin-dependent traction in invasive advance, a key role for microtubules in maintaining elongated, directional migration, and a residual traction-poor invasion component that persists when myosin II activity is inhibited.

## Results

### GBM neurospheres spontaneously invade Matrigel with elongated, protrusion-rich morphologies

Patient-derived GBM neurospheres isolated from human glioblastoma tissue were embedded in Matrigel, a basement membrane–like extracellular matrix, to model 3D invasion. Within the first hours after embedding, neurospheres and cells at the periphery appeared largely rounded. Over time, GBM cells established front–rear polarity and invaded the surrounding matrix, forming elongated cell bodies and pseudopodial protrusions consistent with mesenchymal-like 3D migration (**Figure 1A**). Similar elongated, spindle-like morphologies have been reported for perivascularly invading glioblastoma cells *in vivo*^14-16^ and were also observed in our GBM–iPSC-derived forebrain organoid co-culture model^17-20^ (**Figure S1**), supporting Matrigel-embedded neurospheres as a relevant and tractable platform to study GBM invasion.

**Figure 1.**
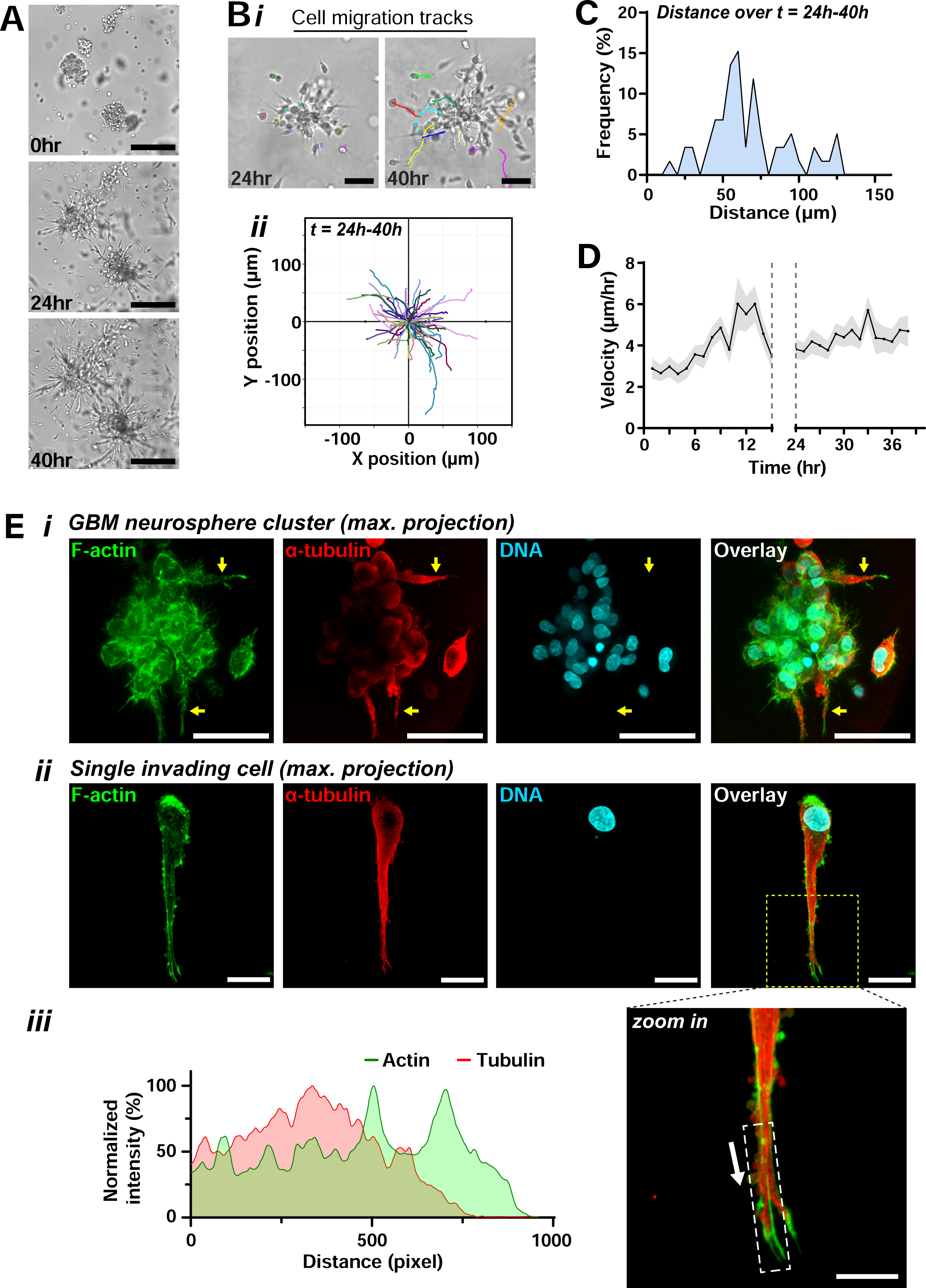
Invasive phenotype of patient-derived GBM neurospheres. **(A)** Brightfield images of GBM neurospheres embedded in Matrigel show pronounced cellular invasion. The invading cells exhibit an elongated morphology with prominent leading-edge protrusions, characteristic of 3D mesenchymal migration. Scale bar = 250 µm. **(B) (i)** Representative images of cell migration tracks analyzed between the 24-hour and 40-hour timepoints. Scale bar = 100 µm. **(ii)** Cell trajectories plotted using x and y coordinates normalized to a single origin, showing displacements of individual cells tracked for a total of 16 hours. Each color represents an individual cell (n = 60 cells from 3 experiments). **(C)** Frequency distribution of migration distances quantified from the tracks in Figure 1B. **(D)** Average velocity of invading cells over time, with an increase during the first 8 hours followed by steady motility (mean±SEM; n = 60 cells from 3 experiments). **(E) (i)** Maximum-intensity projection of confocal sections taken from a GBM neurosphere cluster stained for F-actin (green), α-tubulin (red), and DNA (cyan), revealing the invading protrusions that are enriched for both F-actin and microtubule (yellow arrows). Scale bar = 50 µm. **(ii)** Similar enrichment of F-actin and microtubule was also observed in the singular invading cells. Scale bar = 20 µm. **(iii)** Zoomed-in image from the yellow-dash box of Figure 1Eii and intensity profile showing the extension of F-actin beyond the microtubule. The white arrow indicates the direction of the intensity profile and the white dash box indicates the area used to derive the intensity profile. Scale bar = 10 µm.

To quantify invasion dynamics, brightfield time-lapse imaging of the neurospheres embedded in Matrigel was performed at 30-minute intervals for two days (**Video 1**). The movement of individual cells was tracked over time and trajectory analysis showed predominantly linear paths (**Figure 1B**), suggesting a level of persistence and directionality to the invasion. Within the highly migratory time points of 24-40 hours, the average migration distance was 69.12 ± 3.88 µm (mean ± SEM, **Figure 1C**). Additionally, from the overall migration velocity quantification (**Figure 1D**), the cells invasion speed increased during the first ∼8 hours of tracking and then stabilized, indicating an early acceleration phase followed by sustained motility.

Cytoskeletal organization during invasion was assessed by high-resolution confocal imaging. Invasive protrusions showed dense F-actin enrichment at cellular periphery and prominent microtubules extending within the protrusion shaft (**Figure 1E i–ii**). Fluorescence intensity profiles in single invading cells indicated that F-actin extended beyond the distal ends of microtubules at the leading edge (**Figure 1E iii**). This pattern is consistent with actin-driven protrusion, with microtubules supporting protrusion elongation and polarized migration. Similar dendritic, pseudopodial protrusions have been reported for cells migrating in soft 3D matrices, where microtubules contribute to persistent elongation and guidance^21-23^.

Together, these data establish a robust GBM invasion model in Matrigel characterized by elongated, protrusion-rich migration supported by coordinated actin and microtubule architecture.

### 3D traction force microscopy reveals progressive traction accumulation and force hotspots at protrusion tips

Given the prominent protrusions and their likely mechanical coupling to the surrounding matrix, we used 3D traction force microscopy (3D TFM) to quantify cell–matrix interactions. GBM neurospheres were labeled with SPY-Actin, a non-toxic, cell-permeable live-cell F-actin probe, and embedded in Matrigel containing 0.2-µm fluorescent beads. Confocal time-lapse imaging over 48 h captured actin dynamics together with bead displacements (**Figure 2A, Video 2**). Bead displacements were used to reconstruct the cumulative matrix deformation vector field using the open-source software Saenopy^24,25^ (**Figure 2B**). As defined in Methods, the cumulative deformation field reports the total matrix deformation accumulated up to each time point. Protrusion formation coincided with localized matrix deformations surrounding protrusion tips, indicating robust mechanical interaction between invading GBM cells and the Matrigel. Over time, the cumulative matrix deformation increased as invasion progressed, consistent with the progressive clustering of fluorescent beads around the neurospheres (**Figures 2A and S2**), indicative of substantial and persistent matrix remodeling.

**Figure 2.**
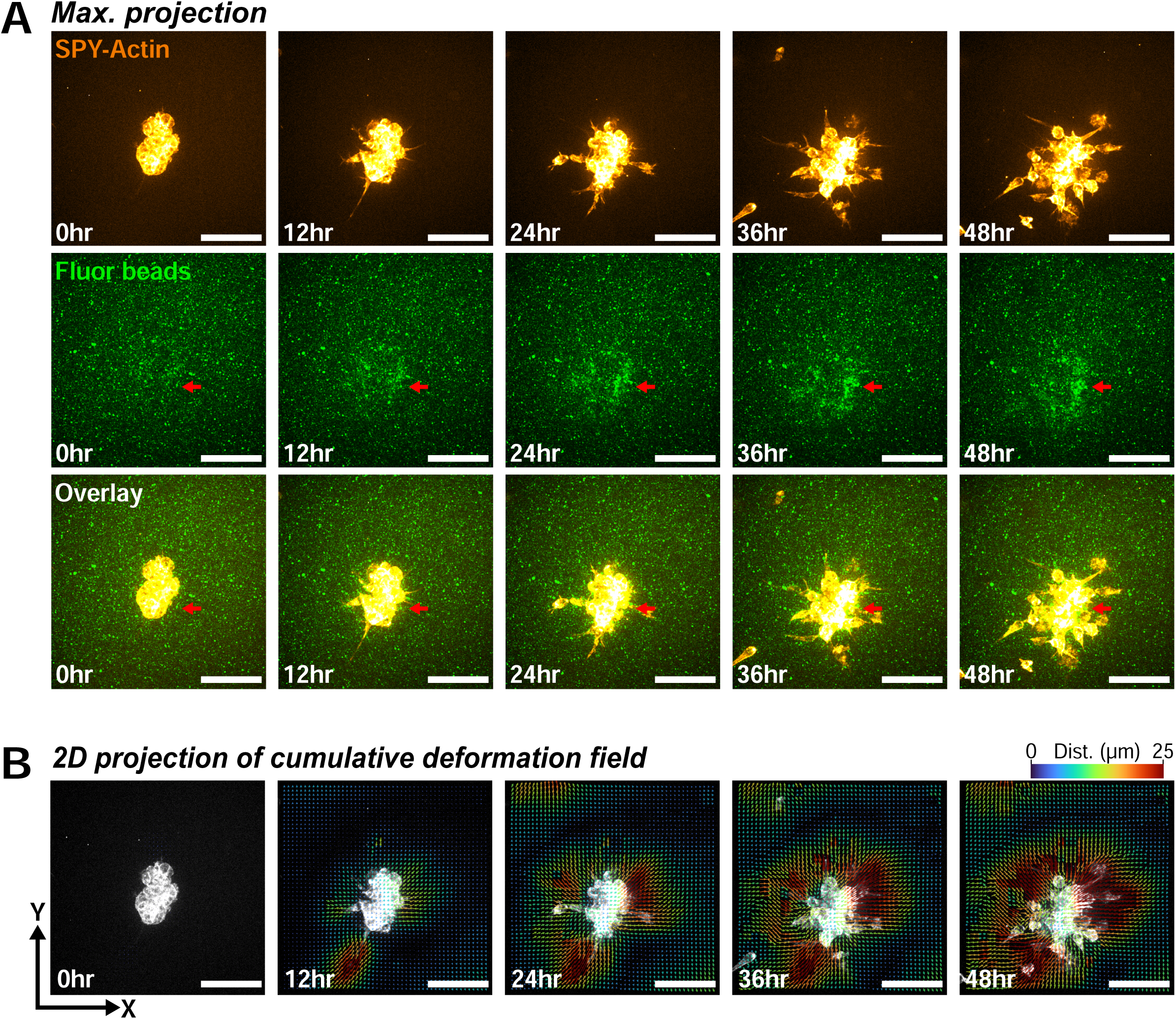
Traction force microscopy (TFM) reveals high-level interactions between the invading GBM cells and their microenvironment. **(A)** Maximum-intensity projections from live confocal imaging of a GBM neurosphere stained with SPY-Actin and embedded in Matrigel supplemented with 0.2 µm fluorescent beads. Actin imaging was used to monitor cell invasion, while bead displacement over time reports cell-matrix interactions. Accumulation of beads around the neurospheres (red arrow) indicates sustained and high-level mechanical engagement between invading cells and the surrounding Matrigel. Scale bar = 100 µm. **(B)** Two-dimensional (2D) projection of the cumulative deformation vector field derived from bead displacement over time. The projection represents the summed displacement across all z-planes. “Cumulative” indicates that the analysis incorporates the total accumulated displacement over the entire imaging period. Arrowheads indicate the direction of bead movement, and the arrow length and color denote the magnitude of the displacement. Scale bar = 100 µm.

Saenopy was then used to convert the interpolated cumulative deformation fields into reconstructed traction force fields generated by the invading GBM cells (**Figure 3A**). At the 24 hr time point, high traction forces were localized around active protrusions, consistent with the deformation patterns observed in Figure 2B. To quantify the overall mechanical output of each neurosphere, we summed the traction forces within the imaging volume at each time point to obtain the total traction force over time (**Figure 3B**). Total traction force increased in a biphasic manner: an initial phase of rapid increase during the first ∼20 hr, followed by a second phase with a slower, gradual increase through the end of imaging. This time-dependent pattern indicates that traction generation evolves dynamically during neurosphere invasion.

**Figure 3.**
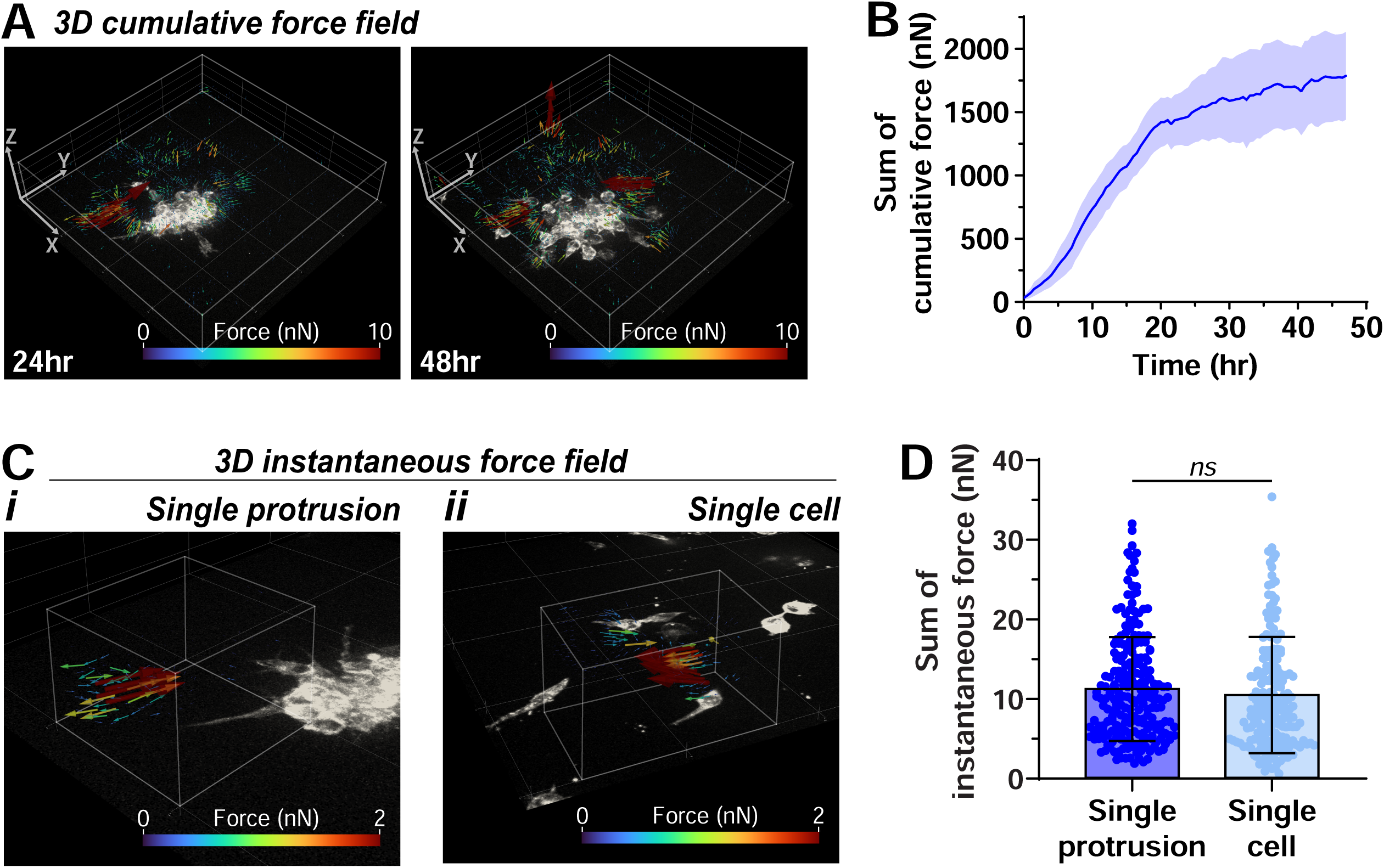
Traction forces increase over time and show similar instantaneous magnitudes in cluster and single cell protrusions. **(A)** Three-dimensional (3D) vector field of the cumulative traction forces derived from the GBM neurosphere in Figure 2. Arrowheads indicate the direction of the traction force, and the arrow length and color denote the magnitude of the force. The white bounding box indicates the region considered in the TFM analysis. Maximum-intensity projections of SPY-Actin staining indicate the positions of the GBM cells. **(B)** The traction force values plotted over time represent the sum of cumulative force within the analyzed region. The data show a progressive increase in traction force, with a rapid rise during the first ∼20 hours followed by a slower, continued increase thereafter (mean±SEM; n = 3 neurospheres from 3 experiments). **(C)** 3D vector fields of the instantaneous traction forces generated by **(i)** a single protrusion from a GBM neurosphere and **(ii)** a single invading GBM cell. ‘Instantaneous’ refers to forces calculated from bead displacements occurring between successive imaging time points (every 30 minutes). The white bounding box marks the region used for the TFM analysis. Maximum-intensity projections of SPY-Actin staining indicate the positions of the GBM cells. **(D)** Sum of instantaneous traction forces generated by single protrusions from GBM neurospheres and by single invading GBM cells, showing no significant difference between the two groups (mean±SD; neurosphere protrusions: n = 13, quantified over 2–47 time points, 3 neurospheres from 3 experiments; single cells: n = 10, quantified over 6–34 time points from 6 experiments).

The slower second phase coincided with the period when cells begin to detach from the neurosphere and invade the matrix (**Video 2**), raising the question of whether early neurosphere protrusions generate higher traction than protrusions from individual invading cells. To directly compare these scenarios, we computed instantaneous traction force fields for selected protrusions, i.e. forces inferred from deformations between consecutive time points (see Methods), and quantified the total force associated with each protrusion (**Figure 3C**). The traction forces generated by neurosphere protrusions were comparable to those generated by protrusions from single invading GBM cells (**Figure 3D**), suggesting that detachment and single-cell invasion do not require higher protrusion traction forces than those produced by already-migrating individual cells.

### Perturbing actin, microtubules, or myosin II differentially impairs invasion and traction generation

Actin polymerization, microtubule dynamics, and myosin II activity jointly govern protrusion formation, polarity, and force transmission during cell migration^26-29^. To assess their contributions to GBM invasion, neurospheres embedded in Matrigel were treated with blebbistatin (myosin II inhibitor)^30^, colchicine (microtubule polymerization inhibitor)^31^, or latrunculin A (actin polymerization inhibitor)^32^. Drug concentrations were selected based on prior studies^33-35^.

As expected, vehicle-treated controls (DMSO) formed elongated protrusions and invaded robustly into Matrigel (**Figure 4A**, **Video 3**), comparable to untreated neurospheres in Figure 1. Inhibiting myosin II with blebbistatin markedly reduced invasion, as reflected by shorter trajectories (**Figure 4B**) and a significant decrease in both migration distance and velocity relative to controls (**Figure 4C–D**). However, blebbistatin-treated cells still produced elongated protrusions and maintained an elongated morphology, supported by aspect ratio measurements (**Figure 4E**), and their trajectories remained largely linear. Together, these findings indicate that myosin II activity is required for efficient invasion but is less critical for maintaining cellular polarization and migration directionality.

**Figure 4.**
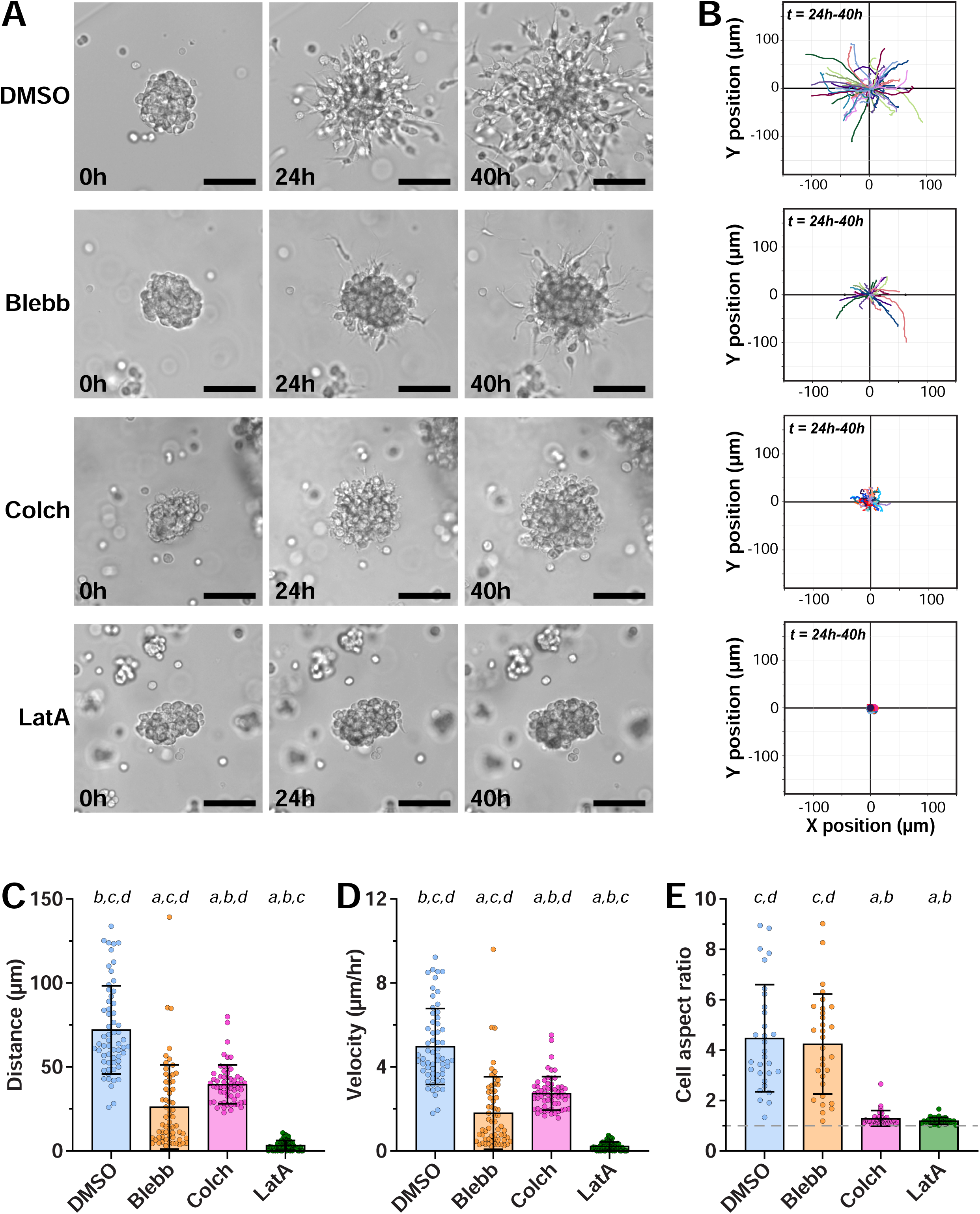
Cytoskeletal perturbations differentially affect GBM neurosphere invasion in 3D Matrigel. **(A)** Bright-field images of GBM neurospheres embedded in Matrigel and imaged over time following treatment with small molecule inhibitors targeting actomyosin contractility (blebbistatin), microtubule polymerization (colchicine), and actin polymerization (latrunculin A). All cytoskeletal inhibitors reduce invasion relative to DMSO, with latrunculin A fully blocking GBM invasion. Scale bar = 100 µm. **(B)** Cell migration tracks quantified between 24–40 hours for each treatment condition. **(C)** Migration distance measurements showing decreased displacement across all inhibitor treatments compared with DMSO. Blebbistatin and colchicine treated cells retain partial migration, whereas latrunculin A treated cells show no detectable migration (mean±SD, n = 60 cells from 3 experiments, *^a^p*<0.001 vs. DMSO, *^b^p*<0.001 vs. blebbistatin, *^c^p*<0.001 vs. colchicine, *^d^p*<0.001 vs. latrunculin A). **(D)** Migration velocity quantification, demonstrating trends consistent with the migration distance results in panel (C) (mean±SD, n = 60 cells from 3 experiments, *^a^p*<0.001 vs. DMSO, *^b^p*<0.001 vs. blebbistatin, *^c^p*<0.001 vs. colchicine, *^d^p*<0.001 vs. latrunculin A). **(E)** Cell aspect ratio analysis. Blebbistatin-treated migrating cells remain elongated, whereas colchicine treatment disrupts elongation, indicating that microtubules are required to support the extended protrusions that maintain cell shape and polarity during invasion (mean±SD, n = 30 cells from 3 experiments, *^a^p*<0.001 vs. DMSO, *^b^p*<0.001 vs. blebbistatin, *^c^p*<0.001 vs. colchicine, *^d^p*<0.001 vs. latrunculin A).

Disrupting either actin or microtubule polymerization shifted cells toward a more rounded morphology (**Figure 4E**). Latrunculin A completely abolished invasion (**Figure 4C–D**), suggesting the essential role for F-actin in GBM protrusion formation and invasive motility. In contrast, colchicine-treated cells retained measurable migration distance and velocity, but with reduced directional persistence, as indicated by the less linear trajectories (**Figures 4B and S3**). This suggests that microtubule polymerization is essential for polarity maintenance and directional persistence, rather than being strictly required for motility per se.

To further characterize invasion under blebbistatin and colchicine, we applied 3D TFM to quantify cell–matrix interactions. Under blebbistatin treatment, live F-actin imaging showed limited protrusion formation and only occasional single-cell invasion (**Figure 5Ai, Video 4**). Bead displacements were minimal throughout the imaging period, consistent with the near-zero cumulative deformation field. Accordingly, total traction forces generated by blebbistatin-treated neurospheres were markedly lower than those of untreated neurospheres (**Figure 5Aii–iii**). These data indicate that myosin II activity is required for robust traction force generation, consistent with its role in focal adhesion maturation^36^ and cellular contractility in migrating cells^37^. At the same time, a small amount of protrusion and invasion persisted despite the lack of measurable traction forces, suggesting a residual invasion mode independent to the myosin II–mediated force transmission.

**Figure 5.**
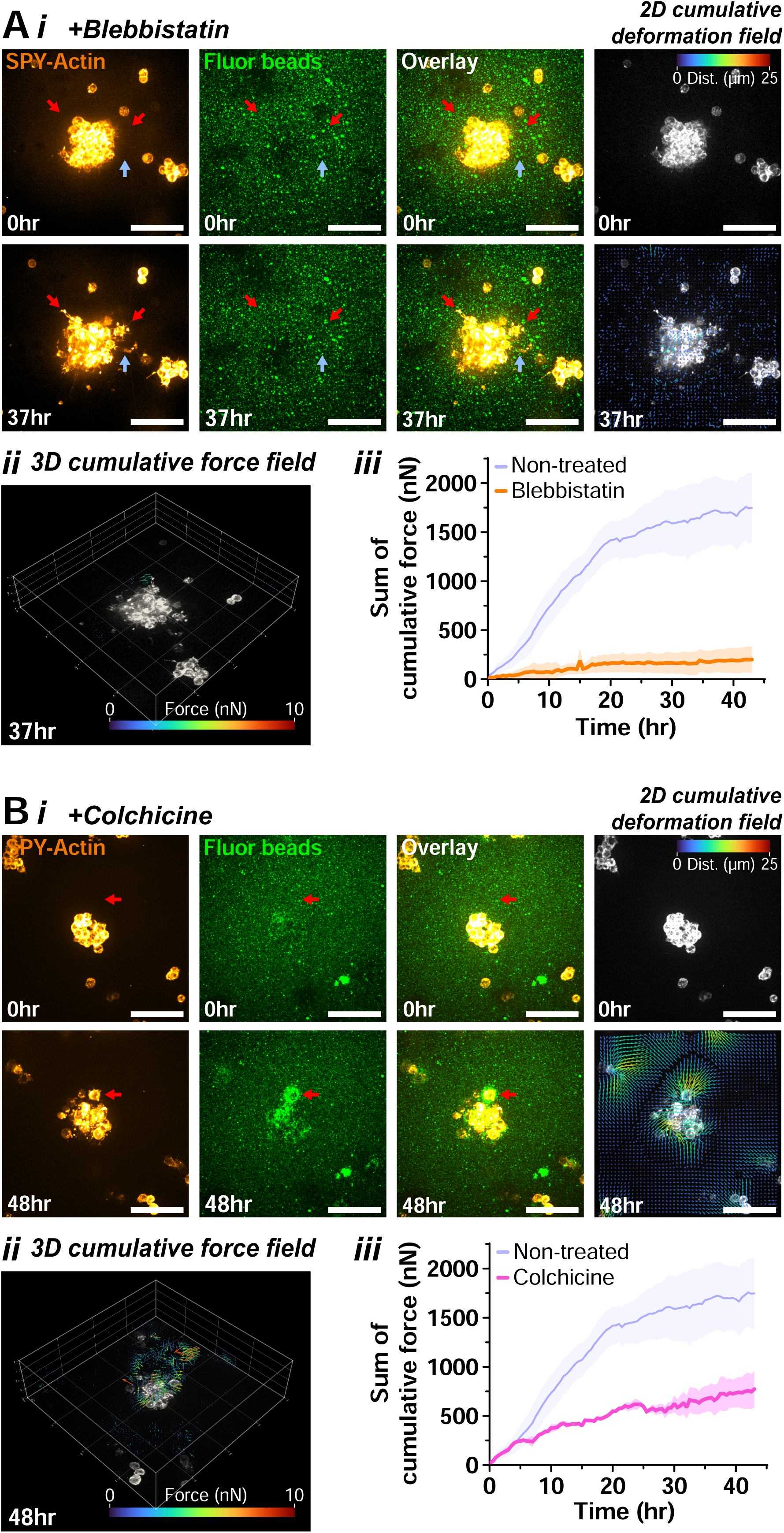
Myosin and microtubule perturbations differentially alter traction force generation during GBM neurosphere invasion. **(A) (i)** Confocal time-lapse images of GBM neurospheres embedded in Matrigel supplemented with 0.2-μm fluorescent beads and treated with blebbistatin. Images represent maximum-intensity projections at 0 hr and 37 hr. SPY-Actin staining highlights minimal protrusive activity, with only limited extensions emerging at later time points (red arrows) and occasional single-cell invasion (blue arrow). Corresponding bead images show negligible bead displacement over time. The 2D projection of the cumulative deformation vector field further demonstrates markedly reduced matrix deformation relative to the untreated neurospheres in Figure 2B. Scale bar = 100 µm. **(ii)** 3D vector field of cumulative traction forces generated by the neurosphere. **(iii)** Sum of cumulative traction forces over time, revealing substantially lower force generation compared with untreated neurospheres (mean±SEM, n = 3 neurospheres from 3 experiments). **(B) (i)** Confocal time-lapse images of GBM neurospheres treated with colchicine under the same imaging conditions as Figure 5A. Maximum-intensity bead projections show clear clustering and accumulation of beads around the GBM cells (red arrow), consistent with detectable local matrix deformation observed in the 2D cumulative deformation vector field. Scale bar = 100 µm. **(ii)** 3D vector field of cumulative traction forces generated by the neurosphere. **(iii)** Sum of cumulative traction forces over time, showing force levels lower than untreated cells but steadily increasing, indicating that GBM cells retain the ability to mechanically engage and deform the matrix even in the absence of prominent protrusive structures (mean±SEM, n = 3 neurospheres from 3 experiments).

One plausible explanation is that protease-dependent matrix remodeling could reduce steric confinement by enlarging local paths in the matrix, which can lessen the mechanical requirement for myosin II–driven contractility during 3D movement^38,39^. Therefore, we tested whether metalloproteinase (MMP) activity contributes to the invasion observed under myosin II inhibition. The pan-MMP inhibitor, GM6001, did not suppress neurosphere invasion (**Figure S4**), indicating that GBM invasion in Matrigel is not MMP-dependent. In addition, combining GM6001 with blebbistatin did not eliminate the residual invasion, which remained comparable to blebbistatin alone. Thus, the residual invasion observed under myosin II inhibition also does not require MMP activity. In line with our observation, the lack of MMP dependence was also reported for breast cancer cell invasion in Matrigel^40^.

Under colchicine treatment, live imaging revealed multiple short F-actin protrusions around invading cells (**Figure 5Bi, Video 5**). This was accompanied by prominent bead displacement and matrix deformation near the cell periphery. Colchicine-treated neurospheres generated total traction forces in the hundreds of nanonewton (nN) range, which is lower than untreated neurospheres (**Figure 5Bii–iii**). Together, these results show that GBM cells can still exert substantial traction forces in the absence of microtubule polymerization, but force output is reduced. This reduction may reflect the impaired formation of long, directed protrusions and the decreased directional persistence.

## Discussion

This study provides a mechanical view of patient-derived GBM neurosphere invasion in Matrigel by combining live imaging, 3D TFM, and targeted cytoskeletal perturbations. Invading GBM cells adopted an elongated protrusive morphology, with F-actin enriched at the cell periphery and microtubules extending along the protrusion shaft, consistent with a mesenchymal-like invasion program. 3D TFM further revealed sustained matrix engagement during invasion, with increasing cumulative traction forces and traction hotspots concentrated near protrusion tips. Pharmacologic perturbations separated cytoskeletal requirements for invasion and force transmission: actin polymerization was essential for invasion, myosin II activity was required for robust traction generation and efficient invasion, and microtubule polymerization supported directional persistence and maximal traction output. Notably, a low level of invasion persisted under myosin II inhibition despite minimal detectable traction forces, and this residual invasion was not suppressed by pan-MMP inhibition. Together, these data identify a traction-poor invasion component that is MMP-independent in this Matrigel model.

Several aspects of these conclusions should be interpreted with some limitations. For example, the invasion behavior described here may be specific to GBM neurospheres cultured in Matrigel, a basement membrane–rich hydrogel composed primarily of laminin and collagen IV. GBM cells may therefore display different invasion phenotypes in other commonly used 3D matrices, including collagen I^41,42^, hyaluronic acid^43^, alginate^44,45^, or polyethylene glycol hydrogels^46,47^. In addition, the brain parenchyma is often better conceptualized as a cell-dense environment with relatively sparse fibrillar ECM. *In vivo*, GBM cells often migrate along perivascular spaces and through cell-dense brain tissue^14^. This differs from the ECM-rich environment modeled by many hydrogels. Despite these differences, the protrusion-rich, mesenchymal-like invasion phenotype observed in our Matrigel culture is consistent with GBM invasion behaviors reported *in vivo*^14-16^, in ex vivo brain slice models^35,48-50^, and in GBM–brain organoid co-culture systems^17-20^, supporting Matrigel-embedded neurospheres as a relevant and tractable platform for quantitative mechanobiology measurements. Looking forward, granular microgel platforms may provide a useful intermediate model by approximating cell-scale crowding while retaining experimental control; microgels can be functionalized with cell–cell junction proteins or ECM ligands to tune the local environment^51-53^.

Our findings also relate to emerging efforts to therapeutically target the mechanical machinery of GBM invasion. The importance of non-muscle myosin II in GBM invasion has been reported in multiple studies^10-13^. Invasive GBM cells often extend elongated protrusions at the leading edge, followed by actomyosin-driven contraction that advances the cell body and retracts the rear^10,12^. Indeed, myosin IIA inhibition suppresses invasion in established GBM models^13^, motivating strategies to inhibit myosin-dependent motility. A recent study described a highly specific myosin II inhibitor, MT-125, with strong brain penetrance and a favorable safety profile^54^. The authors reported that MT-125 suppresses GBM invasion and cytokinesis while increasing redox stress through disruption of mitochondrial fission. MT-125 is now being evaluated in a Phase I clinical trial (NCT07185880). In this context, our data adds an important consideration. Although myosin II inhibition markedly reduced traction forces and strongly impaired invasion, a small population of cells retained invasive behavior despite minimal detectable traction. This suggests an invasion component that is less dependent on myosin II–mediated force transmission. Consequently, myosin II inhibition alone may not fully abolish GBM invasion. Mechanistic dissection of this traction-poor invasion may identify complementary targets that could be combined with myosin II inhibitors such as MT-125 to achieve more complete suppression of invasive spread.

Beyond therapeutic implications, recent work has highlighted the potential value of in vitro invasion characterization for predicting patient prognosis^55^. In this context, traction force measurements during invasion could provide an additional parameter to strengthen such prediction efforts. A practical advantage of our approach is accessibility, as it combines commercially available, ready-to-use Matrigel with Saenopy, a publicly available 3D TFM analysis platform. This should facilitate adoption by other laboratories and support future translational workflows. Nevertheless, larger studies across more patient-derived samples will be required to determine whether traction-based metrics robustly capture GBM invasion phenotypes and add predictive value for clinical outcomes.

Finally, these data contribute to a broader discussion about the diversity of mesenchymal migration programs. The mesenchymal invasion phenotype observed for GBM has been reported previously and is supported by multiple studies^14,16,56^. Increasingly, however, “mesenchymal migration” is recognized as a heterogeneous set of behaviors rather than a single uniform program, where cells can adopt distinct 3D mesenchymal migration mechanisms depending on matrix architecture, confinement, and cell-intrinsic state^57^. Consistent with this view, a recent study showed that fibroblasts and mesenchymal cancer cells can share a 3D migration phenotype driven by traction forces and ECM deformation, but the strength and dynamics of this phenotype are cell-type dependent, highlighting separable sub-phenotypes within mesenchymal migration^58^. Differences can also emerge within a single cancer lineage. For example, breast cancer cell line MDA-MB-231 derivatives selected from different metastatic sites (brain and bone) displayed differential invasion-associated behaviors relative to each other and to the parental line, suggesting that metastatic adaptation can shift where cells fall along the mesenchymal migration spectrum^58,59^. Together, these observations support the concept of mesenchymal migration subtypes that differ in their reliance on actin-rich protrusions, myosin-driven contractility, and traction-mediated matrix engagement. Within this context, our traction-resolved measurements provide a quantitative parameter to position patient-derived GBM invasion along this spectrum, and standardized traction metrics may serve as mechanistic descriptors for comparing mesenchymal subtypes across cell types and microenvironments.

## Materials and methods

### Cell culture

Patient-derived GBM neurospheres, HCM-BROD-0195-C71 (ATCC PDM-140), were generated by the Human Cancer Models Initiative (HCMI) and acquired from ATCC. Neurospheres were cultured in complete medium containing NeuroCult NS-A Basal Medium (StemCell Technologies #05750) supplemented with NS-A Proliferation Supplement (StemCell Technologies #05753), 20 ng/mL EGF (Peprotech #AF-100-15), 20 ng/mL bFGF (StemCell Technologies #78003.1), and 2 µg/mL Heparin (StemCell Technologies #07980) in non-adherent 6-well plates (Corning #3814). Maintenance and cryopreservation followed ATCC guidelines.

For drug inhibitor experiments, GBM neurospheres were embedded in Matrigel (Corning). Briefly, neurospheres were suspended in ice-cold Matrigel, and 15 µL of the suspension (∼15,000 cells) was dispensed into each well of a pre-warmed (∼37°C) 48-well plate. Plates were then incubated at 37°C for 15 min to allow the Matrigel to polymerize. Drug inhibitors, including Blebbistatin (100 µM, MilliporeSigma), Colchicine (25 µM, Fisher Scientific), Latrunculin A (1 µM, Fisher Scientific) and pan-MMP inhibitor GM 6001 (20 µM, MilliporeSigma), were diluted in complete GBM medium and added to the wells. The concentration of DMSO vehicle control matched that of the inhibitor with the highest concentration. Neurospheres were treated for 3 days, with growth and invasion potential monitored daily via z-stack imaging on an Olympus IX71 microscope (10X/0.3NA). All image analyses were performed using ImageJ^60^.

### Live-cell imaging and cell tracking

GBM neurospheres were cultured in Matrigel and seeded in 48-well plates under different treatment conditions (drug inhibitors). Live-cell imaging was performed using an Olympus IX71 microscope with a 10X/0.3NA objective, acquiring images for 15 hours at 30-minute intervals per day over 3 consecutive days. During image pre-processing, five regions of interest (ROIs) of equal size were selected for each condition for cell tracking analysis. The MTrackJ plugin was used to manually track cell movement between 24 and 40 hours (16-hour interval) for each condition. The use of this plugin follows the referenced paper^61^ and the online manual. Cell trajectories were plotted in Graphpad Prism 10 using the x and y coordinates extracted from ImageJ. Given these measurements, cell migration distance and velocity of invading cells were calculated and quantified.

### 3D Traction force microscopy (TFM) analysis

For the experimental set-up, Matrigel was mixed with 0.2 µm Fluospheres carboxylate fluorescent beads (Thermo Scientific #F8810, red fluorescent 580/605) at a 1:600 dilution. Beads were vortexed for 1 minute and thoroughly mixed by pipetting before dilution to ensure uniform distribution and bead density. GBM neurospheres were seeded at 15,000 cells per 15µL Matrigel domes on an 8-well Lab-Tek glass chambered coverglass (Thermo Scientific #155409PK). To track migrated cells, neurospheres embedded in Matrigel domes were stained with SPY555-Actin (Cytoskeleton) at a 1:4000 dilution in complete GBM medium for 2-3 hours before live-cell imaging. All live-cell imaging were performed using Andor Revolution Spinning Disk Laser Confocal Microscope with a 20X/0.75NA objective, while the samples were inside a 37°C 5% CO_2_ imaging incubation chamber.

GBM neurosphere traction forces are quantified from time-lapse image stacks using the open-source program Saenopy^24^. Image stacks of fluorescent beads embedded in Matrigel with a voxel size of 0.69 µm x 0.69 µm x 2 µm and a time delta of 30 min between consecutive recordings were loaded in the analysis pipeline of Saenopy, and deformations between consecutive stacks were determined by particle image velocimetry using the following parameters: window size of 20 pixels, element size of 15 pixels, signal-to-noise threshold of 1.3, and the drift correction was applied. The resulting 3D deformation field were then interpolated to a volume corresponding to the image stack size, with the mesh element size of 7 µm. Two options are available for this interpolation step: (1) The ‘Cumulative’ option locally accumulates deformations over time at each voxel, generating a cumulative deformation field that visualizes the total deformation up to a given time point. (2) The ‘Next’ option calculates deformations between consecutive image stacks at each time step, generating an instantaneous deformation field that visualizes the deformation between a given time point and the preceding one.

The interpolated deformations fields were used by the iterative force reconstruction algorithm to derive the 3D force fields that best describe the observed deformations at each point in time. The material parameters for Matrigel were obtained by rotational rheology as k_0_=675 Pa (linear stiffness), d_0_=1.0 (buckling coefficient), d_s_=0.275 (strain stiffening coefficient), and λ_s_=0.175 (strain stiffening onset). From the reconstructed 3D force fields, we computed the neurosphere traction forces by projecting the individual force vectors towards the epicenter of the deformation fields and summing up the projected force vectors.

### Immunofluorescence staining

Hydrogels were fixed in 2% paraformaldehyde (VWR) and 1% glutaraldehyde (VWR) for 20 min, permeabilized with 0.5% Triton X-100 (MilliporeSigma) for 10 min, and blocked with 5% bovine serum albumin (VWR) for 30 min. Samples were incubated overnight at 4°C with primary antibody against α-tubulin (SigmaAldrich #T9026, DM1A mouse monoclonal) and then with corresponding secondary antibody for 2 hours at room temperature. Nuclei and F-actin were stained with 2 µg/mL DAPI and phalloidin-tetramethylrhodamine B isothiocyanate (Sigma-Aldrich) for 1 hour at room temperature. Images were acquired as z-stacks using Zeiss LSM 880 Confocal Microscope (Plan-Apochromat 63x/1.40 Oil DIC M27).

### Lentiviral transduction on GBM neurospheres

To obtain concentrated lentivirus, HEK293T cells were plated at a starting density of 3×10^6^ cells/mL in a 10 cm culture dish containing culture media (DMEM with sodium pyruvate, 10% FBS, 1% Pen/Strep) approximately 18-24 hours prior to transfection and incubated at 37°C in 5% CO2. Transfections were performed following the TransIT-Lentivirus System protocol (Mirus Bio Catalog #MIR6604) with minor modifications. Briefly, lentiviral packaging plasmids VSV-G (envelope vector), psPAX2 (second-generation lentiviral packaging vector), and the plasmid of interest (pLenti Lifeact-mRuby2 BlastR was a gift from Ghassan Mouneimne^62^, Addgene plasmid # 84384 ; http://n2t.net/addgene:84384 ; RRID:Addgene_84384) were combined in a single tube at a 1:3:4 ratio of VSV-G:psPAX2:plasmid of interest, using 1 µg DNA to 3 µL TransIT reagent in Opti-MEM reduced-serum medium. Transfection complexes were incubated at room temperature for 10 minutes.

Following incubation, the culture medium of HEK293T cells were replaced with DMEM supplemented with 10% FBS and no Pen/Strep. Transfection complexes were added dropwise to the cells, which were then incubated for 48 hours. Lentiviral particles were harvested by collecting the culture medium and centrifuging at 200 × g for 5 minutes to separate cellular debris. The viral supernatant was transferred to a new 15mL conical tube and concentrated using Lenti-X concentrator (Takara Bio, Catalog #631231) by adding one-third volume of concentrator to the volume of supernatant, followed by overnight incubation at 4°C. The next day, concentrated virus was centrifuged at 1,500 × g for 45 minutes at 4°C. The resulting viral pellet was resuspended in 1 mL of complete GBM medium and aliquoted into 0.25 mL volumes in 1.5 mL tubes. For long-term storage, viral aliquots were snap-frozen in liquid nitrogen and stored at -80°C.

For lentiviral transduction of GBM neurospheres in suspension, neurospheres were seeded in 6-well ULA plate at a density of about 500,000 cells/mL and incubated at 37°C in 5% CO_2_ overnight. The next day, the medium was replaced with lentiviral transduction medium consisting of 250 µL concentrated lentivirus supplemented with 1 µL Polybrene in complete GBM medium. Twenty-four hours post-transduction, neurosphere-virus mixtures were transferred to 5 mL tubes and centrifuged at 200 × g for 5 minutes. The resulting cell pellets were resuspended in complete GBM medium and cultured for up to 1 week for recovery, with medium changes every 2-3 days. Successfully transduced neurospheres were selected for 8 days using appropriate antibiotics (2 µg/mL Puromycin or 10 µg/mL Blasticidin).

### Forebrain organoids and GBM neurospheres fusion experiments

Forebrain organoids were differentiated from human induced pluripotent stem cells (iPSCs). The human iPSCs were established from human foreskin fibroblasts that were transfected with plasmid DNA encoding reprogramming factors octamer-binding transcription factor 4 (OCT4), NANOG, SRY-box transcription factor 2 (SOX2), and LIN28. The human iPSCs were maintained in mTeSR Plus serum free medium (StemCell Technologies) on growth factor-reduced Matrigel-coated surface (Corning). The cells were passaged every five to seven days using Accutase and seeded at 1×10^6^ cells per well of a six-well plate in the presence of rho-associated protein kinase (ROCK) inhibitor Y27632 (10 μM, Sigma-Aldrich) for the first 24 hours^63,64^. For the forebrain organoid differentiation, 3x10^6^ iPSCs were seeded in ultra-low attachment (ULA) 24-well plates in 1 mL DMEM/F-12 supplemented with 2% B-27 and 10 µM ROCK inhibitor Y-27632 and incubated overnight. Differentiation was carried out for 15 days by changing the medium every other day with DMEM/F-12 supplemented with 10 µM SB-431542 (ALK5 inhibitor) and 100 nM LDN-193189 (ALK2 inhibitor). Differentiated forebrain organoids were subsequently cultured in a 24-well ULA plate for the duration of the experiment, with 50% medium changes every other day using forebrain organoid complete media (DMEM/F-12, 2% B-27, 1% Pen/Strep). We received the differentiated forebrain organoids on day 15 and proceeded with fusion experiments for 16-day and 20-day old organoids. Using wide-bore pipette tips, individual forebrain organoids were transferred into 96-well V-bottom plates in complete GBM medium were GBM neurospheres were seeded at around 50,000 cells per well in a final volume of 200 µL. Forebrain organoids and GBM neurospheres were co-cultured for 24 hours at 37°C in 5% CO_2_ incubator. The following day, fused organoids were transferred using wide-bore pipette tips into 24-well ULA plates and cultured in complete GBM medium for the duration of the experiment. Growth and successful fusion of organoids and GBM neurospheres were monitored over time through imaging using Olympus microscope (10X).

### Statistical analysis

All graphs and analyses were generated using Graphpad Prism 10. Figure legends specify the sample size for each condition. Statistical analyses were conducted using paired t-test and one-way analysis of variance (ANOVA) with Tukey multiple comparison test. All statistical tests were considered significant if *p* < 0.05.

## Acknowledgements

The authors would like to thank the Confocal Microscopy Laboratory at FSU College of Medicine and Prof. Yuan Wang for access to the confocal microscopes. The authors would like to thank Dr. Terra Bradley for the careful editing of the manuscript. The authors would like to thank all lab members in Dr. Jerome Irianto’s and Dr. Yue Julia Wang’s lab for their support and valuable suggestions in reviewing the manuscript. HS, JI, and BS were supported by startup funds from Florida State University, an award from the Institute of Pediatric Rare Diseases (FSU College of Medicine), and awards from the Florida Department of Health’s Bankhead-Coley Cancer Research Program (award number 21B11), and Live Like Bella Pediatric Cancer Research Initiative (award number 23L06). YL and FS were supported by NSF award #2425703. DB was supported by the German Research Foundation (DFG; project 326998133 – TRR-SFB 225 – subprojects A01). The GBM neurospheres are from the Human Cancer Models Initiative (HCMI), https://ocg.cancer.gov/programs/HCMI.

---

**Figure S1.**
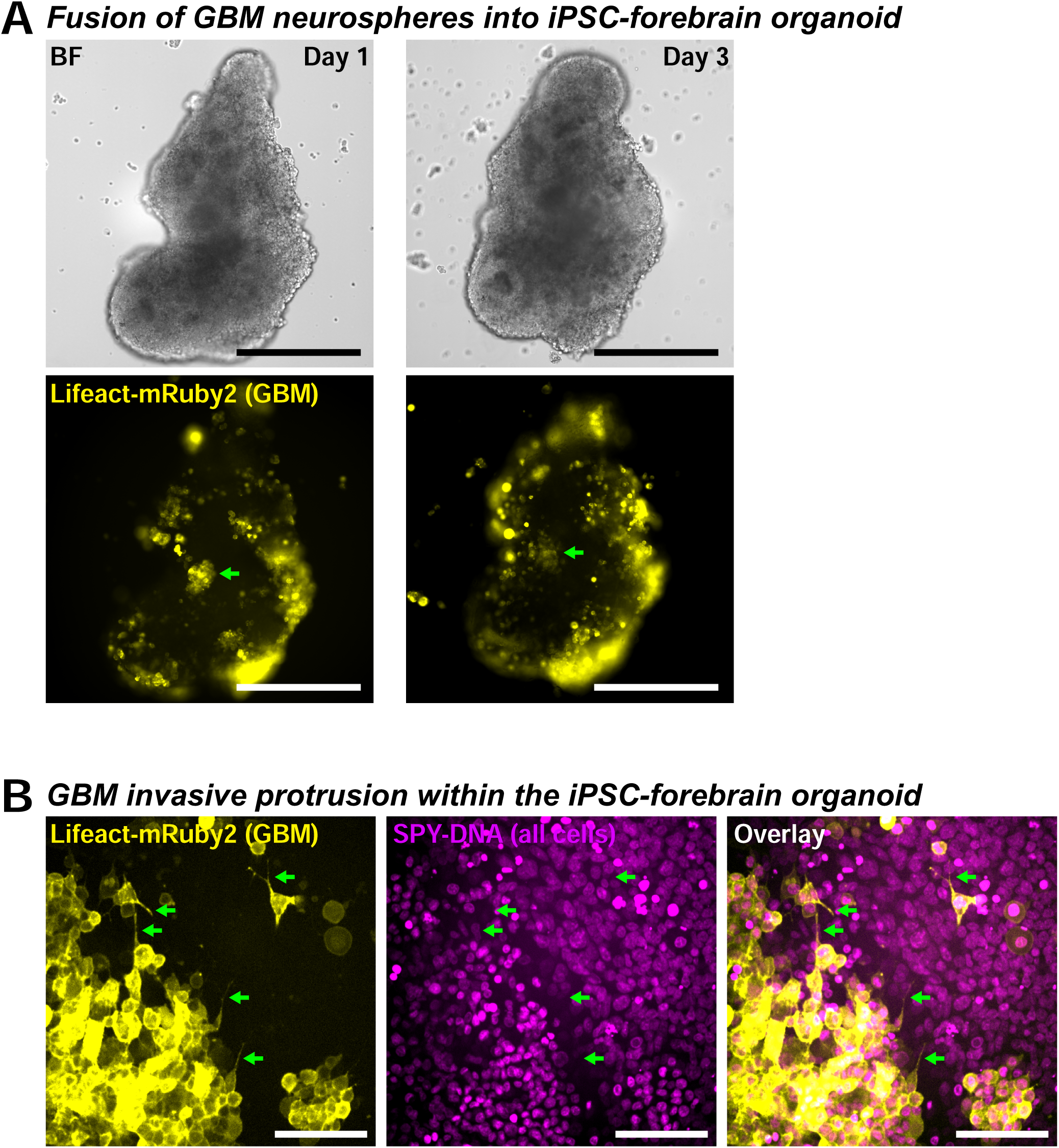
Invasion of GBM cells into the induced pluripotent stem cells (iPSC) derived forebrain organoids. **(A)** GBM neurospheres expressing the actin filament–binding protein Lifeact–mRuby2 were fused with iPSC-derived forebrain organoids. GBM cells were incorporated into the organoids one day after fusion. By day 3, the GBM cells had invaded the forebrain organoids (green arrow). Scale bar = 500 µm. **(B)** Invading GBM cells at day 3 displayed extensive protrusions (green arrows), similar to those observed during GBM invasion in Matrigel (Figure 1E). SPY-DNA was used to label the nuclei of both GBM cells and the forebrain organoid. Scale bar = 100 µm.

**Figure S2.**
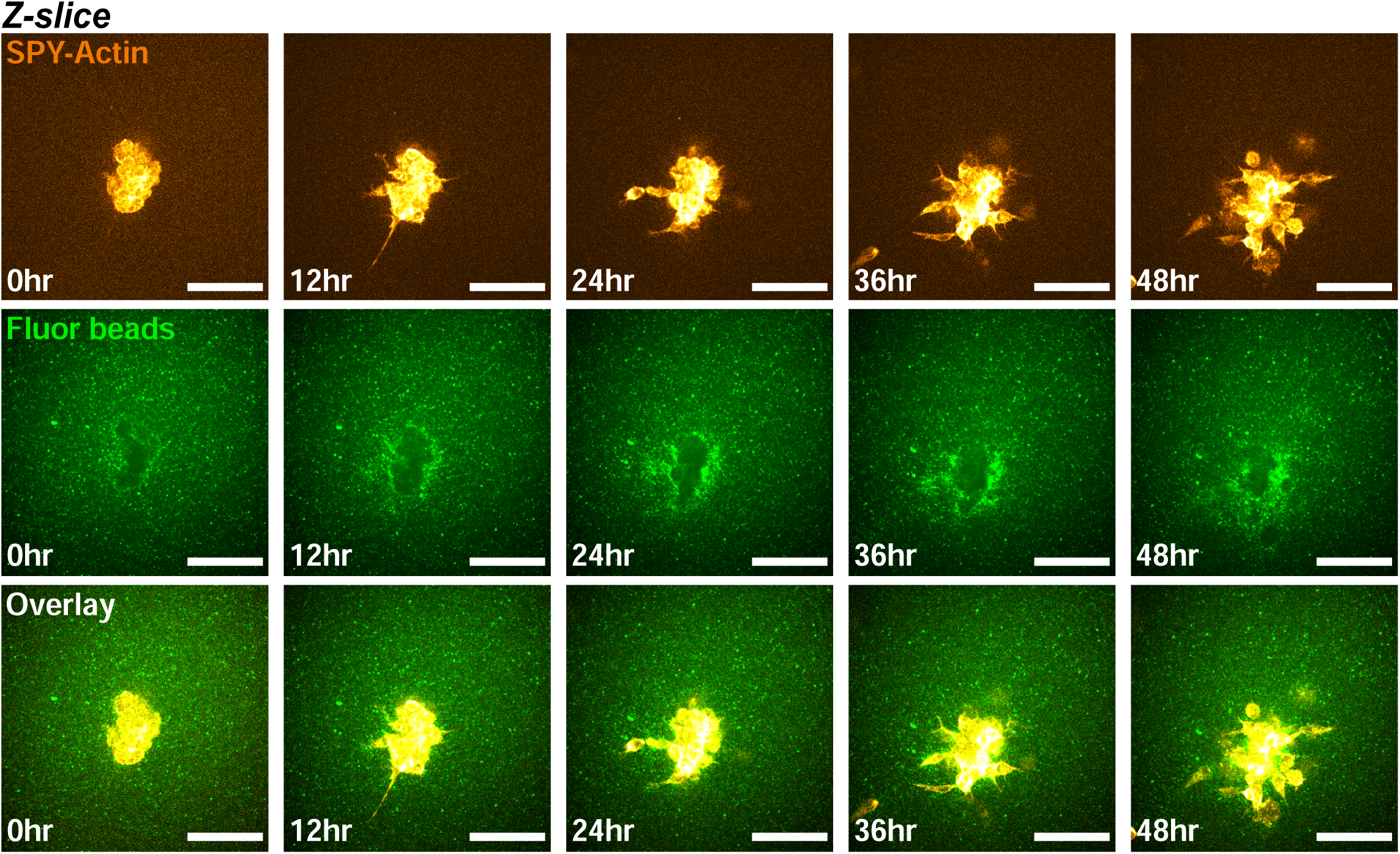
Sustained cell-ECM interaction during GBM invasion. A representative confocal section from the live imaging shown in Figure 2A. Clustering of fluorescent beads around the GBM neurosphere prior to invasion into the surrounding Matrigel suggests strong and sustained interactions between the cells and the extracellular matrix.

**Figure S3.**
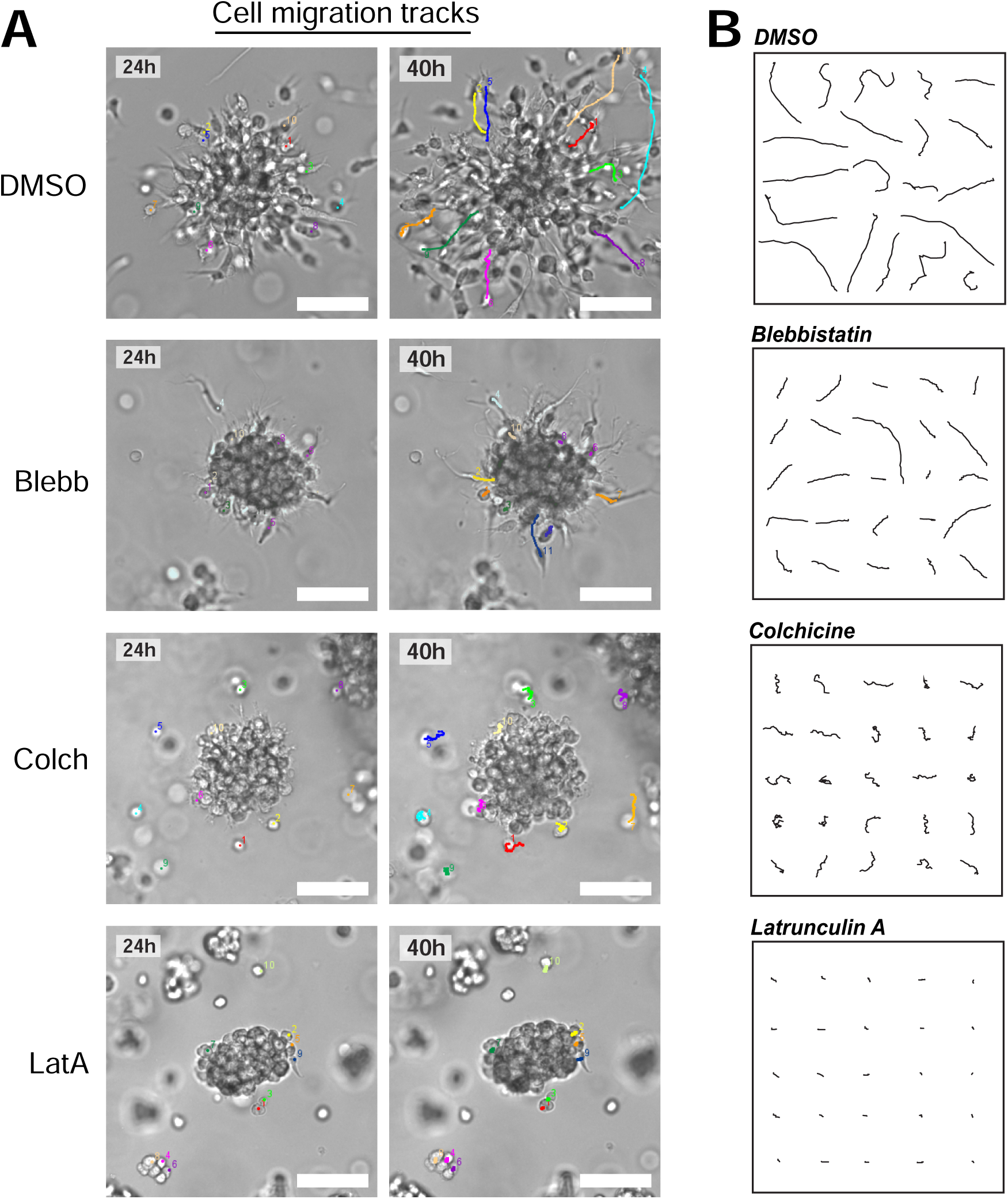
Effect of cytoskeletal inhibitors in GBM invasion. **(A)** Representative cell migration tracks using MTrackJ (ImageJ). GBM neurospheres embedded in Matrigel were treated with blebbistatin, colchicine, and latrunculin A. Migration of cells were tracked for a total of 16 hours between the 24-hour and 40-hour timeframe in all conditions. Each individual cell is color-coded and numbered accordingly. Scale bar = 100 µm. **(B)** Visual comparison of 20-25 randomly selected cells from each condition, highlighting migration directionality.

**Figure S4.**
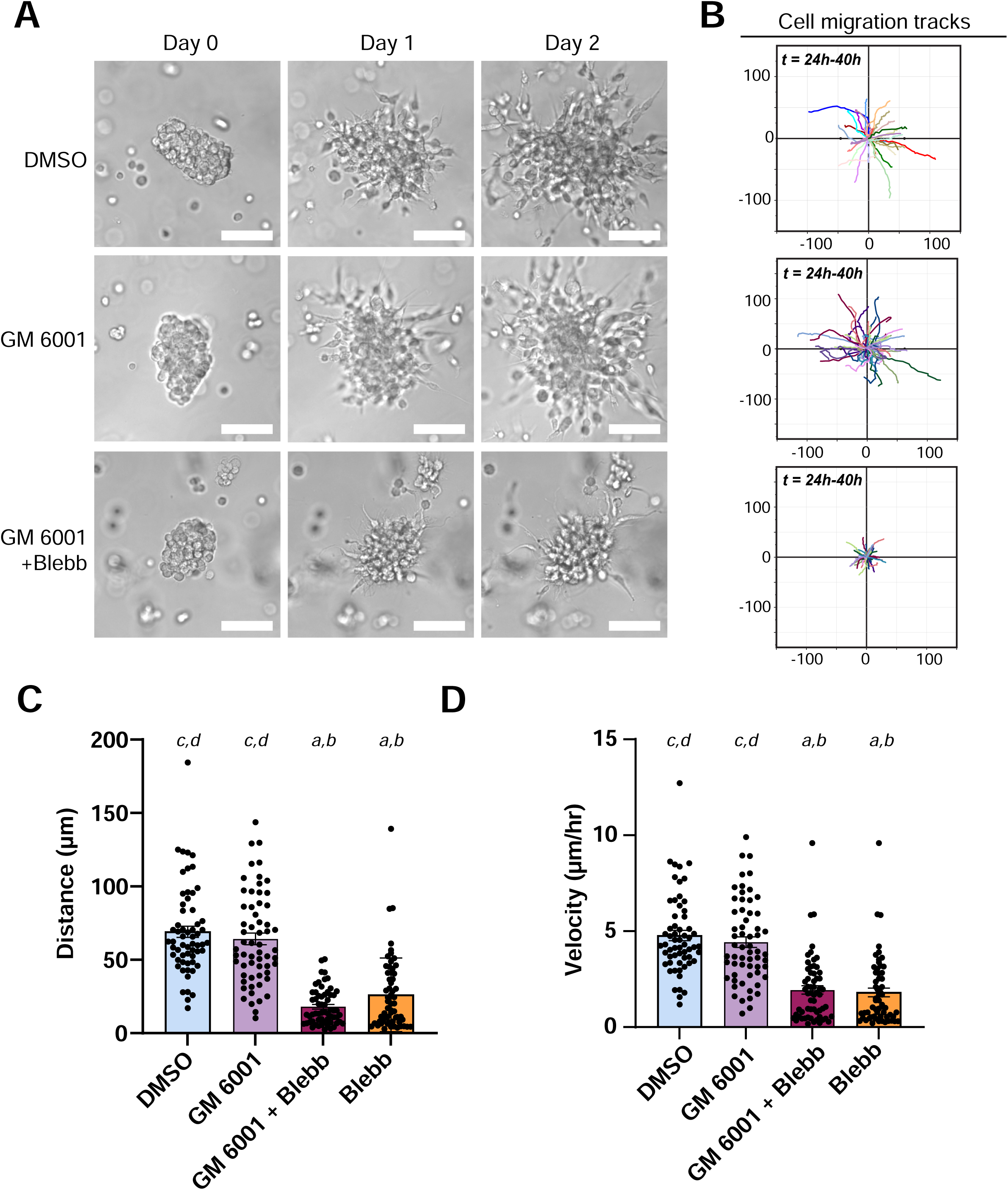
Metalloproteinase (MMP) inhibition does not suppress GBM invasion in Matrigel. **(A)** Representative bright-field images of GBM neurospheres treated with DMSO, the pan-MMP inhibitor GM6001, or GM6001 combined with the myosin II inhibitor blebbistatin. Inhibition of MMP activity alone does not reduce GBM invasion. Furthermore, the limited invasion observed following myosin inhibition persists in the presence of GM6001, suggesting that the residual invasion under myosin inhibition is not dependent on MMP activity. Scale bar = 100 μm. **(B)** Cell migration tracks quantified between 24–40 hours for each treatment condition. **(C)** Quantification of migration distance showing that GM6001 treatment does not significantly alter GBM invasion compared with DMSO controls. In addition, GM6001 does not further reduce migration when combined with blebbistatin (mean±SEM, n = 60 cells from 3 experiments, *^a^p*<0.001 vs. DMSO, *^b^p*<0.001 vs. blebbistatin, *^c^p*<0.001 vs. colchicine, *^d^p*<0.001 vs. latrunculin A). **(D)** Migration velocity quantification showing trends consistent with the migration distance measurements in panel (C) (mean±SEM, n = 60 cells from 3 experiments, *^a^p*<0.001 vs. DMSO, *^b^p*<0.001 vs. blebbistatin, *^c^p*<0.001 vs. colchicine, *^d^p*<0.001 vs. latrunculin A).

## Video legends

**Video 1**

Brightfield live imaging reveals the invasion of GBM neurospheres when cultured in Matrigel. The imaging was performed at 0-16hr and 24-40hr after seeding.

**Video 2**

Confocal time-lapse imaging of a GBM neurosphere stained with SPY-Actin and embedded in Matrigel supplemented with 0.2-µm fluorescent beads. The video shows the maximum-intensity projection of the actin-labeled cells and the fluorescent beads, alongside the corresponding 2D cumulative deformation field derived from bead displacement and the summed cumulative traction forces over time.

**Video 3**

Brightfield live imaging of representative GBM neurospheres treated with DMSO (control), blebbistatin, colchicine, and latrunculin A. The imaging was performed at 0-16hr and 24-40hr after seeding.

**Video 4**

Confocal time-lapse imaging of a GBM neurosphere stained with SPY-Actin and embedded in Matrigel supplemented with 0.2-µm fluorescent beads following treatment with the myosin II inhibitor blebbistatin. The corresponding 2D cumulative deformation field shows minimal bead displacement compared with the untreated neurosphere shown in Video 2.

**Video 5**

Confocal time-lapse imaging of a GBM neurosphere stained with SPY-Actin and embedded in Matrigel supplemented with 0.2-µm fluorescent beads following treatment with the microtubule polymerization inhibitor colchicine. The corresponding 2D cumulative deformation field shows moderate bead displacement compared with the untreated neurosphere shown in Video 2.

